# *Sincell:* Bioconductor package for the statistical assessment of cell-state hierarchies from single-cell RNA-seq data

**DOI:** 10.1101/014472

**Authors:** Miguel Juliá, Amalio Telenti, Antonio Rausell

## Abstract

**Summary:** Cell differentiation processes are achieved through a continuum of hierarchical intermediate cell-states that might be captured by single-cell RNA seq. Existing computational approaches for the assessment of cell-state hierarchies from single-cell data might be formalized under a general framework composed of i) a metric to assess cell-to-cell similarities (combined or not with a dimensionality reduction step), and ii) a graph-building algorithm (optionally making use of a cells-clustering step). *Sincell* R package implements a methodological toolbox allowing flexible workflows under such framework. Furthermore, *Sincell* contributes new algorithms to provide cell-state hierarchies with statistical support while accounting for stochastic factors in single-cell RNA seq. Graphical representations and functional association tests are provided to interpret hierarchies. *Sincell* functionalities are illustrated in a real case study where its ability to discriminate noisy from stable cell-state hierarchies is demonstrated.

**Availability and implementation:** *Sincell* is an open-source R/Bioconductor package available at http://bioconductor.org/packages/3.1/bioc/html/sincell.html. A detailed manual and vignette describing functions and workflows is provided with the package.

## 1 INTRODUCTION

Unbiased profiling of individual cells through single-cell RNA-seq allows assessing heterogeneity of transcriptional states within a cell population (Wu *et al.*, 2014). In the context of a cell differentiation or activation process, such transcriptional heterogeneity might reflect a continuum of intermediate cell-states and lineages resulting from dynamic regulatory programs (Qiu *et al.*, 2011; Trapnell *et al.*, 2014; Bendall *et al.*, 2014). Such continuum might be captured through the computational assessment of cell-state hierarchies, where each individual cell is placed in a relative ordering in the transcriptional landscape. Additionally, statistical support should be provided in order to discriminate reliable hierarchies from stochastic heterogeneity, arising from both technical (Brennecke *et al.*, 2013) and biological factors (Raj *et al.*, 2006; Rand *et* al., 2012; Shalek *et* al., 2013; McDavid *et al.*, 2014; Deng *et al.*, 2014).

A number of algorithms have been used to assess cell-state hierarchies from single-cell data (Qiu *et al.*, 2011; Bendall *et al.*, 2014; Amir *et al.*, 2013; Trapnell *et al.*, 2014; Jaitin *et al.*, 2014). These approaches might be formalized under a general framework (Supplementary Table S1). Here we present *Sincell*, an R/Bioconductor package where the various building blocks of that general workflow are extended and combined (**Figure 1**). Notably, *Sincell* implements algorithms to provide statistical support to the cell-state hierarchies derived from single-cell RNA-seq. The package is complemented with graphical representations and functional association tests to help interpreting the results.

## 2 DESCRIPTION

### 2.1 Integrative framework for assessment of cell-state hierarchies

As input, *Sincell* requires an expression matrix with user-defined normalized gene expression levels per each singlecell in the study (**Figure 1**). First, a cell-to-cell distance matrix is calculated through a metric of choice. *Sincell* provides both linear and non-linear distances: Euclidean, Mutual Information, L1 distance, Pearson and Spearman correlation. Optionally, the distance matrix may be obtained from the dimensions lead by a dimensionality reduction algorithm, performed to keep the most informative part of the data while excluding noise. Both linear and non-linear algorithms are provided: Principal Component Analysis (PCA), Independent Component Analysis (ICA), t-Distributed Stochastic Neighbor Embedding (t-SNE) and non-metric Multidimensional Scaling (MDS).

**Figure 1.**
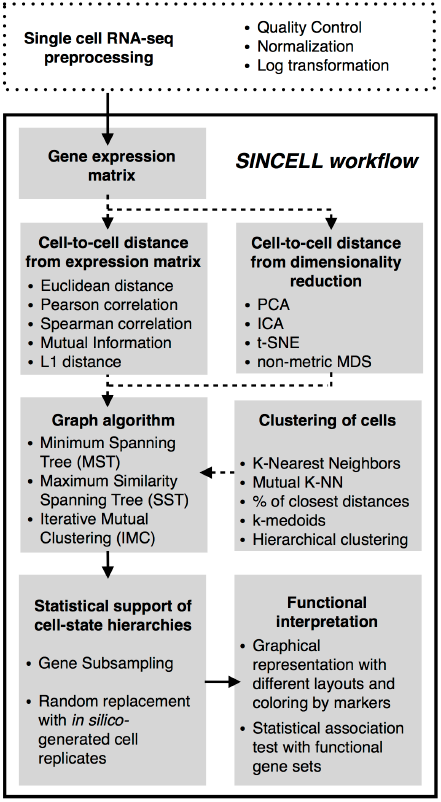
Overall workflow for the statistical assessment of cell-state hierarchies implemented by the *Sincell* R package. Dashed arrows correspond to optional steps in the analysis

Second, a cell-state hierarchy is obtained by applying a graph-building algorithm on the cell-to-cell distance matrix. Graph-building algorithms may consider cells both individually or in clusters of highly similar cells. *Sincell* provides different clustering methods (e.g. K-Mutual Nearest Neighbours, k-medoids, agglomerative clustering, etc.) as well as graph-building algorithms (MST, SST and IMC; **Figure 1** and **Supplementary Text**).

Stochastic factors -both technical and biological-may drive cell-state heterogeneity observed on Single-cell RNA seq data. Additionally, hierarchies derived from experiments with a low number of individual cells (e.g. 96 cells when using a Fluidigm C1™ Single-Cell Auto Prep System) are more susceptible to noise artifacts than experiments profiling thousands of individual cells (e.g. flow cytometry data). *Sincell* implements two algorithms to discriminate reliable hierarchies from noise-driven ones. The first strategy relies on a gene resampling procedure. The second one is based on random cell substitution with *in silico*-generated cell replicates. These replicates are built by perturbing observed gene expression levels with random noise, following patterns of stochasticity described in single-cell RNA-seq (Brennecke *et al.*, 2013; Anders and Huber, 2010; Shalek *et al.*, 2014). Either approach generates a population of hierarchies whose similarities to the reference one show the distribution of the hierarchy stability against changes in the data. Details of the algorithms are described in the **Supplementary Text**.

### 2.2 Graphical representations and functional association tests for interpreting cell-state hierarchies

*Sincell* provides graphical representations of cell-to-cell similarities in low-dimensional space as well as graph displays of cell-state hierarchies. The possibility of coloring cells by expression levels of a gene of choice helps inspecting the agreement of the hierarchy with selected cell markers. Furthermore, *Sincell* implements an algorithm to determine the statistical significance of the association of the hierarchy with the expression levels of a given gene set (**Supplementary Text**). Gene lists defined by molecular signatures, Gene Ontology terms or by pathway databases can be systematically evaluated.

## 3 APPLICATION

*Sincell* R package is accompanied with a detailed vignette illustrating all previous functionalities in real single-cell RNA-seq data. We use data from (Trapnell *et al.*, 2014) quantifying gene expression levels in differentiating myoblast at 0, 24, 48 and 72 hours. The original report describes a continuum in the differentiation process by building a cell-state hierarchy where individual cells from all time points were taken together. Here we analyze each time-point separately and evaluate the statistical evidence of cell-state heterogeneity within them (**Supplementary Figure 1**). Our results show that early times of differentiation produce unstable hierarchies suggesting a low degree of cell-state heterogeneity. However, late differentiation times produce statistically significant hierarchies that reflect cell-state diversity along the differentiation process.

## 4 DISCUSSION

The landscape of computational approaches to assess cell-state hierarchies from single-cell data is far from being fully explored. The diversity of biological studies and rapid singlecell technological evolution require a comprehensive toolbox where users may easily tailor workflows and compare alternative methods and assumptions. Furthermore, cell-state hierarchies should be statistically supported before being used as input in subsequent analyses. *Sincell* R package addresses these needs by providing a general analysis framework, new algorithms for statistical support as well as tools for functional interpretation of cell-state hierarchies.

*Funding*: Supported by The European FP7 grant number 305762 and Swiss National Science Foundation no. 149724. Part of the computations were performed at the Vital-IT (http://www.vital-it.ch) Center for high-performance computing of the SIB Swiss Institute of Bioinformatics.

*Conflict of interest:* none declared.

## Supplementary Information

**Supplementary Table 1.** Computational methods for the assessment of cell-state hierarchies. The table shows a list of published approaches for the assessment of cell-state heterogeneity from single-cell data together with their main methodological features. The last row includes the standard Principal Component Analysis (PCA) to reflect its use in single-cell data analysis; in this case two references are provided as a non-exhaustive list of examples.

| Method | Reference | Single-cell data | Dimensionality reduction | Metric to assess cell-to-cell distances | Clustering algorithm | Graph -building algorithm / ordering representation | Trajectory assessment | Software availability |
| --- | --- | --- | --- | --- | --- | --- | --- | --- |
| **SPADE** | Qiu et al 2011 | Mass Cytometry Data and Flow Cytometry Data | NA | L1 distance | Agglomerative clustering | Minimum Spanning Tree (MST) | NA | R/Bioconductor |
| **Wanderlust** | Bendall et al 2014 | Mass Cytometry Data | NA | Cosine distance | NA | K-Nearest Neighbours Graph (K-NNG) | A single non-branching trajectory is assessed from an average of "shortest path"- trajectories over an ensemble of l-out-of-k-nearest-neighbor graphs (l-k-NNGs) | Matlab based |
| **viSNE** | Amir et al 2013 | Mass Cytometry Data and Flow Cytometry Data | t-Distributed Stochastic Neighbor Embedding (t-SNE) | Distance in low-dimensional space | NA | NA | NA | Matlab based |
| **Monocle** | Trapnell et al 2014 | Single-cell RNA-seq | Independent Component Analysis (ICA) | Distance in low-dimensional space | NA | Minimum Spanning Tree (MST) | Longest path through MST is used to define branching trajectories and ordering in "pseudo-time" | R/Bioconductor |
| **Jaitin et al 2014** | Jaitin et al 2014 | Single-cell RNA-seq | NA | Correlation | Hierarchical clustering + manual definition of seeds | Circular projection (CAP) of posterior probabilities of association with the model's classes | NA | Not available |
| **PCA** | Dalerba et al. 2011<br>Treutlein et al 2014 | Single-cell RNA-seq | Principal Component Analysis (PCA) | Distance in low-dimensional space | NA | NA | NA | Multiple platforms |

NA: Not Applicable

**Supplementary Figure 1.**
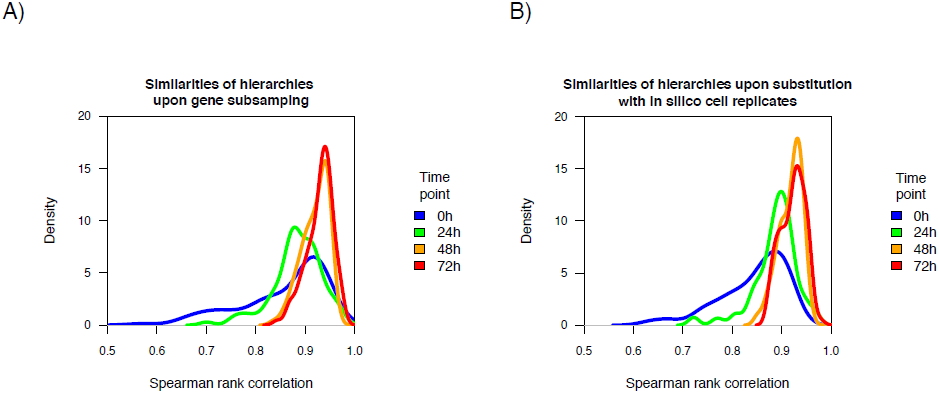
Statistical support for cell-state hierarchies obtained in differentiating myoblast samples at 4 time points (0, 24, 48 and 72h) from Trapnell et al 2014. **A**. Similarities of hierarchies upon random gene subsampling. The figure represents the distribution of similarities between a reference cell-state hierarchy and the 100 hierarchies obtained when 100 random sets of 50% of genes are subsampled. **B**. Similarities of hierarchies upon random cell replacement with *in silico* cell replicates. The figure represents the distribution of similarities between a given cell-state hierarchy and the 100 hierarchies obtained when 100 % of individual cells are substituted by a randomly chosen *in silico* replicate of themselves. One thousand *in silico* replicates were generated for each cell with default parameters. Four distributions are represented in each panel corresponding to the hierarchies obtained at different time points: 0, 24, 48 and 72 hours (blue, green, orange and red respectively). A distribution of similarities with a high median and a low variance is indicative of a cell-state hierarchy robust to variations in the data. See **Supplementary Text** for details.

### Supplementary Text

#### Before starting using *Sincell*

*Sincell* workflow starts from an expression matrix gathering the gene expression levels for every single-cell in the experiment. Before starting using *Sincell*, quality controls to filter out individual cells from the analysis have to be performed by the user. Expression levels need also to be previously normalized to account for library size or technical variability (e.g. through the use of spike-in molecules). Variance stabilization through log-transformation is also recommended.

#### Novel graph-building algorithms presented in *Sincell*

We present here two graph-building algorithms that can be used to infer the progression through a continuum of intermediate cell states: the *Maximum Similarity Spanning Tree* and the *Iterative Mutual Clustering Graph (IMC).* Both algorithms start from a cell-to-cell distance matrix as an input, and compute a connected graph where nodes represent cells and edges represent their kinship as intermediate cell-states. The weight of an edge connecting two cells corresponds to the original distance between them. The algorithms start with all nodes unconnected, treating them as clusters of size 1.

##### Maximum Similarity Spanning Tree (SST)

In a first iteration, the two clusters with the lowest distance are connected forming a cluster of size 2. In a new iteration, distances among clusters are recomputed and a new connection is added between the next two clusters with the lowest distance. A distance between a cluster of size higher than one and another cluster is the lowest distance between any of their constituent cells. The process is repeated until there are no cells unconnected.

In contrast with the *Minimum Spanning Tree* (MST) algorithm (that minimizes the total sum of the weights of any possible spanning tree), the SST algorithm prioritizes the highest similarities between any two groups of cells, proceeding in an agglomerative way that represents intermediate cell states. In some cases, MST and SST can lead to the same graph.

##### Iterative Mutual Clustering Graph (IMC)

In a first iteration, a connection between two clusters A and B is added if A is among the closest *k* nearest clusters of B and B is among the closest *k* nearest clusters of A. This process is iterated until there are no unconnected cells. As for SST, the distance between a cluster of size higher than one and another cluster is the lowest distance between any of their constituent cells.

#### Algorithmic strategies to provide statistical support to cell-state hierarchies from single-cell RNAseq

The fact that a cell-state hierarchy is obtained by using any given algorithm (e.g. MST, SST or IMC) does not necessarily imply that it reflects a true biological scenario of cell activation/differentiation. It might well be that the hierarchy obtained is mainly driven by noise due to either biological or technical factors. The relative contribution of stochastic factors to the observed differences across cells is expected to be higher if cells within a sample are in a homogeneous steady-state. In that case, a low cell-to-cell heterogeneity will lead to cell-state hierarchies very sensitive to small variations in the initial gene expression data. On the other extreme, high levels of cell-to-cell heterogeneity driven by a real granularity in an activation/differentiation process will translate into robust hierarchies that can be reproduced despite stochastic perturbations of the data.

To help discriminating reliable cell-state hierarchies from noisy rearrangements, *Sincell* implements two algorithms: i) a strategy relying on a gene resampling procedure and ii) an algorithm based on random cell substitution with *in silico*-generated cell replicates.

##### A. Gene resampling

This algorithm performs “s” times a random subsampling of a given number “*n*” of genes in the original gene expression matrix. For each subsampling, a new connected graph of cells is computed using the same method as for the hierarchy being tested. In each subsampling, the similarity between the resulting graph and the original one is assessed as the Spearman rank correlation between the two graphs of the shortest distance for all pairs of cells. The distribution of Spearman rank correlation values of all subsamplings can be interpreted as the distribution of similarities between hierarchies that would be obtained from small changes in the data. A distribution with a high median and small variance would indicate a well-supported cell-state hierarchy. On the contrary, a distribution with a low median of similarities and/or a wide variance would indicate a hierarchy very sensitive to changes in the data, and therefore not well statistically supported.

##### B. Random cell substitution with in silico-generated cell replicates

Gene expression levels detected by single-cell RNA seq are subject to stochastic factors both technical and biological. This means that, if it were possible to profile multiple times the same cell in the same cell-state (or, more realistically, a population of individual cells in a highly homogeneous state), the detected expression levels of a gene would randomly fluctuate within a distribution of values. In the ideal scenario where that distribution was known for each gene, individual cell replicates could be produced *in silico*, leading to variations in gene expression levels similar to what would be obtained from *in vivo* replicates. The generation of *in silico* replicates would then permit testing the reproducibility of the cell-state hierarchy upon random replacement of a fraction of the original cells with them.

##### B1. Generation of in silico cell replicates

The distribution of the expression levels of a gene can be described by a measure of variability such as the variance or the coefficient of variation. It is known that the expected variation is dependent on the mean expression values of the gene (Anders and Huber 2010; Brennecke et al 2013). Based on this, we can simulate a stochastic fluctuation of the expression of a gene by perturbing the observed level in a given cell with an error term whose magnitude is consistent with the mean-variance relationship observed in the data. By doing that in all genes from an individual cell C_i_, we can produce an *in silico* replicate of it.

Sincell implements this strategy as follows: first, the mean *m* and variance *v* of all genes in the original gene expression matrix is computed. Genes are assigned to classes according to the deciles of mean they belong to. Next, for a given gene g, a variance *v* is randomly chosen from the set of variances within the class of the gene. Then, a random value drawn from a uniform distribution *U(0,v)* of mean zero and variance *v* is added to the expression value of a gene *g* in a cell *c.* By perturbing in such a way all genes in a reference cell c, we obtain an *in silico* replicate *c’.* Redoing the process *n* times, *n* stochastic replicates are generated for each original cell. Alternatively, a squared coefficient of variation *cv2* can be randomly chosen from the set of coefficient of variation values within the class of the gene. Then, the variance *v* for the uniform distribution is assessed by *v* = (*cv2* × *m*^2^).

Stochasticity in gene expression at the single-cell level has also been described as following a lognormal distribution *log(x)∼N(m,v)* of mean *m* and variance *v* (Bengtsson et al 2005; Raj et al 2006). More recently, Shalek et al 2014 described gene expression variability in single-cell RNA-seq through a log normal distribution with a third parameter *alpha* describing the proportion of cells where transcript expression was detected above a given threshold level. Authors found that the majority of genes in their study (91%) showed distributions well described by the three-parameter model (p < 0.01, goodness of fit test; Shalek et al 2014). Sincell can use this “three parameter” model estimation to generate random perturbations of gene expression levels and produce *in silico* cell replicates accordingly.

##### B2. Random cell substitution with in silico-generated cell replicates

Once cell-replicates have been generated, a *Sincell* algorithm performs *“s”*times a random replacement of a given number *“n”* cells on the original gene expression matrix with a randomly selected set of *in-silico* replicates. For each set of substitutions “s”, a new connected graph of cells is assessed using the same method as for the hierarchy being tested. In each “s”, the similarity between the resulting graph and the original one is assessed as the Spearman rank correlation between the two graphs of the shortest distance for all pairs of cells. The distribution of Spearman rank correlation values of all replacements might be interpreted as the distribution of similarities between hierarchies that would be obtained from stochastic perturbations of a proportion of cells. A distribution with a high median and small variance would indicate a well-supported cell-state hierarchy. On the contrary, a distribution with a low median of similarities and/or a wide variance would indicate a hierarchy very sensitive to changes in the data, and therefore not well statistically supported.

##### C. Application of Sincell algorithms to provide cell-state hierarchies with statistical support on a real single-cell RNA seq data set

We applied *Sincell* algorithms to provide cell-state hierarchies with statistical support on a publicly available single-cell RNA-seq dataset from Trapnell et al 2014. The authors generated single-cell RNA-seq libraries for differentiating myoblasts at 0, 24, 48 and 72 hours. Original data can be accessed at GEO database accession number GSE52529 (ftp://ftp.ncbi.nlm.nih.gov/geo/series/GSE52nnn/GSE52529/suppl/GSE52529_fpkm_matrix.txt.gz). Following Trapnell et al 2014 and the vignette of its associated Bioconductor package *Monocle* (http://www.bioconductor.org/packages/devel/bioc/html/monocle.html), the expression matrix is restricted to 575 genes differentially expressed between cells from time 0 and the ensemble of cells of times 24, 28 and 72 hours of differentiation. Here, we analyze each time-point separately and evaluate the statistical evidence of cell-state heterogeneity within them.

Four cell-state hierarchies were assessed for each time point separately (0, 24, 48 and 72h) on their log-transformed FPKM values using the first two dimensions of a dimensionality reduction with Independent Component Analysis (ICA) and a Minimum Spanning Tree (MST). To evaluate the statistical support of the arrangements obtained, two *Sincell* algorithms were applied: i) a gene resampling procedure and ii) a random cell substitution with *in* silico-generated cell replicates. **Supplementary Figure 1A** represents the distribution of similarities between a reference cell-state hierarchy and the 100 hierarchies obtained when a random set of 50% of genes are subsampled 100 times. **Supplementary Figure 1B** represents the distribution of similarities between a reference cell-state hierarchy and the 100 hierarchies obtained when 100% of the cells are replaced by a randomly chosen *in silico* replicate of themselves 100 times.

In both cases, late time points lead to hierarchies with a high median while early time points had a lower median and a higher variance. Results suggest that at early time points homogeneity of cell states is high, leading to hierarchies more sensitive to perturbations of the data and therefore less statistically supported. However, late time point showed hierarchies more robust to both gene subsampling and replacement with in-silico replicates, reflecting a marked heterogeneity in cell-states. Indeed, a gradient can be observed in both panels (from 0 to 24, 48 and 72h) suggesting that heterogeneity in cell-states increased as a function of time.

#### Functional association tests to help interpreting cell-state hierarchies

Once a cell-state hierarchy has been assessed and its statistical support checked, the next step is interpreting the hierarchy in functional terms. *Sincell* allows different graphical representations that can help interpreting the hierarchies in terms of the features of the samples (e.g. differentiation time) or the expression levels of markers of interest. In this section, we propose an analytical approach to test whether the cell-state hierarchy associates with a given functional gene set, that is: whether the relative similarities among the individual cells in the hierarchy are driven by the expression levels of a subset of genes with a common functional feature.

*Sincell* implements an algorithm to evaluate this association. First, a new cell-state hierarchy is assessed where only the expression levels of the genes in a given functional gene set are considered. Second, the similarity of that hierarchy with the reference hierarchy (the one assessed on the initial gene expression matrix) is calculated. The similarity between the two hierarchies is computed as the Spearman rank correlation between the two graphs of the shortest distance for all pairs of cells. Third, an empirical p-value of the observed similarity between the two hierarchies is provided. The empirical p-value is derived from a distribution of similarities resulting from random samplings of gene sets of the same size.

This *Sincell* algorithm is particularly suited to evaluate associations with gene set collections such as those from the Molecular Signatures Database (MSigDB) of the Broad Institute (http://www.broadinstitute.org/gsea/msigdb/collections.jsp), gene lists representing Gene Ontology terms of functional pathways, and in general, any gene set collections that might be of particular interest for the user.

